# Oxytocin and Vasopressin at Birth Prevent Hypoactivity and Excess Weight Gain in Vole Offspring Delivered by Cesarean Section

**DOI:** 10.64898/2026.05.02.722408

**Authors:** Miranda E. Partie, Katelyn Rogers, W. Tang Watanasriyakul, Sabreen Ahmed, Phineas Delgado, James E. Blevins, Sara M. Freeman, William M. Kenkel

## Abstract

Birth occurs during a sensitive period in brain development wherein hormones facilitate the dramatic shift in physiology that accomplishes the transition to extrauterine homeostasis. The surge in birth signaling hormones is abridged in cases of delivery by cesarean section (CS), which accounts for 32% of all births in the U.S. Epidemiological studies have associated birth via CS with increased risk of obesity in later life. Here, we sought to investigate this association using an experimental preclinical animal model, the prairie vole. Subjects were delivered either via vaginal delivery (VD) or CS and then cross-fostered. CS delivery led to increased body weight across development, which could be prevented with ‘hormone rescue’ of oxytocin (OXT) and arginine vasopressin (AVP), delivered to neonates immediately after CS. This weight gain could not be attributed to differences in birth weight, parenting, food consumption, or thermoregulation; however, CS subjects moved slower than VD subjects, which hormone rescue reversed.

Hormone rescue also reduced adiposity in adulthood among CS subjects. The dopamine system was dysregulated in the caudate/putamen of CS offspring, suggesting a neural mechanism for the decreased locomotion. Hormone rescue of CS neonates restored dopamine synthesis in the caudate/putamen and increased spontaneous locomotor activity. These findings suggest CS can lead to increased weight gain in part through a reduction of locomotion driven by long-lasting changes in striatal dopamine regulation, all of which can be prevented by treating CS neonates with a single peripheral administration of two birth-signaling hormones, OXT and AVP.

## Introduction

Delivery by cesarean section (CS) accounts for 32.3% of births in the United States as of 2023 [1]. CS is usually a medically necessary procedure with a long track record of saving lives; however, in epidemiological studies it is also associated with a number of adverse outcomes for both mother and offspring [2,3]. Among these, one of the most well-established associations is between delivery by CS and heightened risk for obesity in offspring. Recent meta-analyses have estimated delivery by CS to result in a 23-59% increased risk for childhood obesity [2,4,5]. The association between CS and offspring obesity survives correction for various confounding factors, as can be seen in within-family sibling comparison, where young adults delivered by CS have a 64% increased risk of obesity compared to siblings born by vaginal delivery (VD) [6]. After adjustments for: marital status, maternal race, prenatal tobacco smoke exposure, maternal age, maternal body mass index (BMI), hypertensive disorders during pregnancy, gestational diabetes, prenatal antibiotic use, child sex, parity, and birthweight z-score, children delivered via scheduled CS still experience a 77% higher risk of obesity [7]. Neither does breastfeeding explain this association [7].

We view the neurodevelopmental consequences of CS from a neuroendocrine perspective such that in vaginal delivery (VD), birth includes the surge of several ‘birth-signaling’ hormones critical for the successful adaptation to extra-uterine life [8,9]. Plasma levels of all known birth-signaling hormones are lower immediately after CS delivery. Many of these hormones, particularly the two neuropeptides oxytocin (OXT) and arginine vasopressin (AVP), exert lifelong developmental effects when their levels are manipulated in early life [10,11].

In the current study, we primarily focused on offspring weight gain, which we have corroborated in prior experimental work [12]. Here, we sought to prevent CS-induced weight gain for the first time by administering a cocktail of OXT and AVP directly to CS neonates. Throughout development, we then measured behavior, neurobiology, and anatomy. These tests began with several behaviors relevant to thermoregulation in early life (ultrasonic vocalizations (USVs), thermography, and thermotaxis), because thermoregulation is a key adaptation upon entry into the extrauterine environment and also one that could explain enhanced weight gain in CS offspring. We also assessed how foster parents treated CS offspring across several tests (parental behavior, pup retrieval and alloparental responsiveness) to see if differential care could contribute to the outcomes of CS offspring. Across development, we monitored locomotor activity using two approaches (home cage video and the Rodent Activity Detector (RAD) [13]). We also assessed food consumption and adult thermoregulation. Finally, in adulthood, we assessed several morphometric measures of the body along with the OXT, AVP, and dopamine (DA) systems in the brain at the level of both ligand and receptor to attempt to identify CS-induced changes in (neuro)anatomy.

## Materials and Methods

All procedures were done in accordance with the Institutional Animal Care and Use Committee of the University of Delaware. Additional methodological details can be found in the Supplemental Information.

### Subjects

Subjects consisted of laboratory-bred prairie voles (*Microtus ochrogaster*) descended from a wild-caught stock. Breeding pairs, composed of stud males aged 60-240 days and primiparous females aged 60-90 days, were created to generate the subjects utilized in this study. Following weaning at PND-21, same-sex offspring were re-housed into polycarbonate cages in groups of up to four. Animals had ad libitum access to water and two sources of food, high-fiber Purina rabbit chow (Rabbit Diet HF 5326, LabDiet) and standard mouse chow (RMH 3000, LabDiet) with enviro-dry and cotton nestlets provided for nesting material in breeder cages. Animal cages were maintained at room temperature (22°) and under a 14:10 hour light:dark cycle. Unless otherwise noted, all observations took place between 10:00 AM and 3:00 PM.

### Birth Conditions

To generate experimental subjects, pairs were created according to previously published procedures restated here [12,14]. To minimize the contributions of prematurity, we developed a timed mating paradigm to ensure accurate prediction of expected delivery. Pregnant female dams bore litters either via CS or VD approximately 21.5 days after mating. This CS paradigm typically achieved delivery ∼12 hours prior to expected delivery. In the CS condition, surgical delivery occurred via laparotomy following 90 seconds of CO_2_ anesthesia according to the protocol developed by the Forger lab [15], an approach which avoids the confound of pharmacological anesthesia and the resulting neurodevelopmental consequences. CS delivery was thus a terminal procedure so as to avoid the confound of surgery affecting maternal behavior and because CS vole dams do not respond maternally to pups [16]. Following delivery (CS) or discovery within 16 hours of delivery (VD), all pups were treated and then cross-fostered as same-condition litters to parents that had had litters of their own within the previous 3 days. There were three treatment conditions, consisting of: VD pups treated with saline (‘VD-SAL’), CS pups treated with saline (‘CS-SAL’), or CS pups treated with ‘hormone rescue’ (0.1 mg/kg OXT and 0.1 mg/kg AVP, ‘CS-HR’). cCP TH-ir data were obtained from a separate cohort of animals born under the same conditions as stated above, with the exception that CS-HR subjects received 0.1 mg/kg OXT-only. All injections were carried out subcutaneously. For all measures except TH-ir, 17 VD-SAL litters, 15 CS-SAL litters, and 18 CS-HR litters were generated, yielding 75, 67, and 77 pups respectively. Nineteen VD-SAL litters, 9 CS-SAL litters, and 8 CS-HR litters were generated for TH-ir measures. Supplemental TH-ir measures were obtained from 7 VD and 6 CS litters. Sample sizes for each specific measure are included in the results.

### Home Cage Parental Behavior

On PNDs 1 and 4, the behavior of a subset of a litter’s foster parents was observed for 10 minutes. The number of parents / pups within the nest was recorded once per minute. This yielded recordings from 13 VD-SAL litters, 10 CS-SAL litters, and 8 CS-HR litters.

### Ultrasonic Vocalizations

Following Parental Behavior testing on PNDs 1 and 4, two pups from each litter were individually removed from the home nest and placed into a recording chamber for 5 minutes as we have done previously [17,18]. On PNDs 6 and 9, all pups from the litter were tested similarly. This yielded recordings from 40 VD-SAL, 38 CS-SAL, and 40 CS-HR pups on PNDs 1 and 4, and 93 VD-SAL, 79 CS-SAL, and 73 CS-HR pups on PNDs 6 and 9.

### Pup Retrieval

To gauge how effective CS and VD pups were at eliciting care, we examined the latency to retrieve pups. Immediately after ultrasonic vocalizations on PNDs 1 and 4, when pups were returned to the home cage, the parents were collectively scored for latency to retrieve returned pups. With both parents in the nest, a single pup was returned to the opposite corner of the cage. The latency for that pup to be retrieved back to the nest was then scored via video recording. No distinction between sire and dam was made. This yielded recordings from 42 VD-SAL pups, 34 CS-SAL pups, and 41 CS-HR pups.

### Alloparental Response

To further gauge how effective CS and VD pups were at eliciting care, we compared the responsiveness of vetted adult male alloparents to pups from each condition. In this way, we reversed the typical alloparental test by focusing on the contribution of the pup rather than the alloparent. The alloparent’s behavior was then recorded and scored as we have done previously [14,17,18] by a single observer blind to condition. This yielded recordings from 14 VD-SAL litters, 16 CS-SAL litters, and 16 CS-HR litters.

### Thermography

On PND-7, a subset of litters were tested for thermographic analysis as we have done previously [14]. The results consisted of the number of pixels at or above 34°, the number of huddles, and the total perimeter of all huddles. Because some recordings were lost due to limited battery life on recording devices, this yielded recordings from 13 VD-SAL litters, 10 CS-SAL litters, and 8 CS-HR litters.

### Thermocline

On PND-10, whole litters were tested for thermographic analysis as we have done previously [12] with the singular modification of changing the age at testing from PND-8 to PND-10 to ensure greater pup mobility. Testing on the thermocline lasted 20 minutes and subjects’ positions were scored once every 60 seconds. This yielded recordings from 12 VD-SAL litters, 12 CS-SAL litters, and 13 CS-HR litters.

### Diet and food consumption

All subjects were provided with a 1:1 mixture of traditional vole chow (LabDiet Rabbit Diet HF 5326, ‘Standard diet’) and conventional mouse chow (Prolab® RMH 3000). We have previously termed this diet ‘high fat’ [12] though ‘calorie dense’ would be a more accurate description. Compared to traditional vole chow, this diet produces a modest acceleration of body weight gain, however, it should also be noted that a minority of vole colonies routinely provide a similar diet to their animals (personal communications made via the VoleBase open science platform). Food consumption was measured over a 6 hour duration weekly on the basis of each cage and divided by the number of animals in the cage.

### Body temperature

On PND-30, a subset of subjects were injected subcutaneously on the dorsum with a temperature sensor (BioTherm13 passive integrated transponder, BioMark Inc.). From PND-30 to PND-65 (mean = 44.3 days), these subjects were recorded for 1-7 days at a time. This yielded recordings from 24 VD-SAL subjects (14 female, 10 male), 20 CS-SAL subjects (8 female, 12 male), and 22 CS-HR subjects (12 female, 10 male), with an average of 443 measurements for each subject.

### Home cage locomotor activity

From PND-25 to PND-67, a subset of subjects were subjected to 2 hour recordings in the home cage every week. This yielded recordings of 37-61 VD-SAL voles at any given age, 31-56 CS-SAL voles, and 34-57 CS-HR voles. From PND-60 through PND-77, a subset of subjects’ home cages was equipped with automated locomotor activity detectors [13]. This yielded recordings from 8 cages of VD-SAL females, 7 cages of VD-

SAL males, 4 cages of CS-SAL females, 4 cages of CS-SAL males, 4 cages of CS-HR females, and 6 cages of CS-HR males, with an average of 11,920 observations (i.e. 8.28 days) for each cage.

### Micro-CT

On PND-50, a subset of subjects were lightly anesthetized with 2% isoflurane and 3D-imaged with a Brüker SkyScan 1276 (Kontich, Belgium) to assess visceral adiposity. This yielded measures of visceral adiposity from 14 VD-SAL, 12 CS-SAL, and 19 CS-HR subjects.

### Tissue Collection

On PND-77, subjects were anesthetized with isoflurane followed by cervical dislocation. Vole length and final body weight were recorded. The following tissues were collected and weighed: scapular brown adipose tissue (BAT), anterior (inguinal) + posterior (scapular) subcutaneous white adipose tissue (WAT) pads, as well as livers.

### Immunohistochemistry

Fixed brains were immunohistochemically processed according to previously published methods [19], incubating in primary antibodies (either AVP at 1:16,000 dilution, OXT at 1:16,000, or tyrosine hydroxylase (TH) at 1:3500 MilliporeSigma antibodies # AB1565, MAB5296, and IHCR10056, respectively). OXT-ir and AVP-ir analysis yielded 23 VD-SAL, 18 CS-SAL, and 24 CS-HR subjects. TH-ir analysis yielded 32 VD-SAL, 15 CS-SAL, and 14 CS-HR subjects, and supplemental TH-ir analysis yielded 13 VD and 12 CS subjects.

### Autoradiography

Flash frozen brains were autoradiographically processed for according to previously published methods [20–22], using ^125^I-Ornithine Vasotocin Analog, ^125^I-Linear Vasopressin 1a Receptor Antagonist, or ^25^I-RTI (Revvity). This yielded 20 VD-SAL, 14 CS-SAL, and 18 CS-HR subjects.

### Analysis

All statistical analyses were carried out in R. Birth mode was examined using linear mixed effects modeling, repeated over either the individual subject or litter. Given that terminal outcomes were measured at a single time point and exhibited substantial litter-level variance, we analyzed terminal measures (body length and adipose depots) using litter–sex averages. For repeated measures we first conducted omnibus testing using the lmer() function from the *lme4* library [23]. Contrasts (e.g. by age) were then conducted using Welch’s ANOVA with post-hoc testing (i.e. Group comparisons) carried out with the Games-Howell test, which accounts for unequal variances and sample sizes. Where appropriate, we included litter and sex as covariates. For brain measures, we also included anterior-posterior position as a covariate. However, for the exploratory autoradiography analyses, we limited reporting to main effects of group and applied a Benjamini-Hochberg false detection rate correction to adjust p-values for multiple comparisons. Data are reported as means plus/minus standard error, alpha = 0.05, and effect sizes reported as either η2 or Cohen’s *d*.

## Results

Additional results can be found in the Supplemental Information.

### Weights Across Development

We measured various aspects of behavior relevant to energy balance across development (Figure 1A-B). At delivery, VD-SAL litters were heavier (F(2,47) = 7.4, p < 0.002), however, by PND-4, CS-HR pups had caught up to VD-SAL, and by PND-9, CS-SAL had caught up to VD-SAL, while CS-HR weighed significantly more than CS-SAL (p = 0.025). CS-SAL animals were consistently heavier than VD-SAL animals starting on PND-49 and heavier also than CS-HR animals on PND-63 and 77 (F(2,534,45) = 21.56, p < 0.001, Figure 1D). CS-HR was no different than VD-SAL at any point. Even under the assumption of maximal plausible correlation, there was no relationship between weight on PND-1 and weight in adulthood (*r* = 0.17, p = 0.11, Figure 1E).

**Figure 1.**
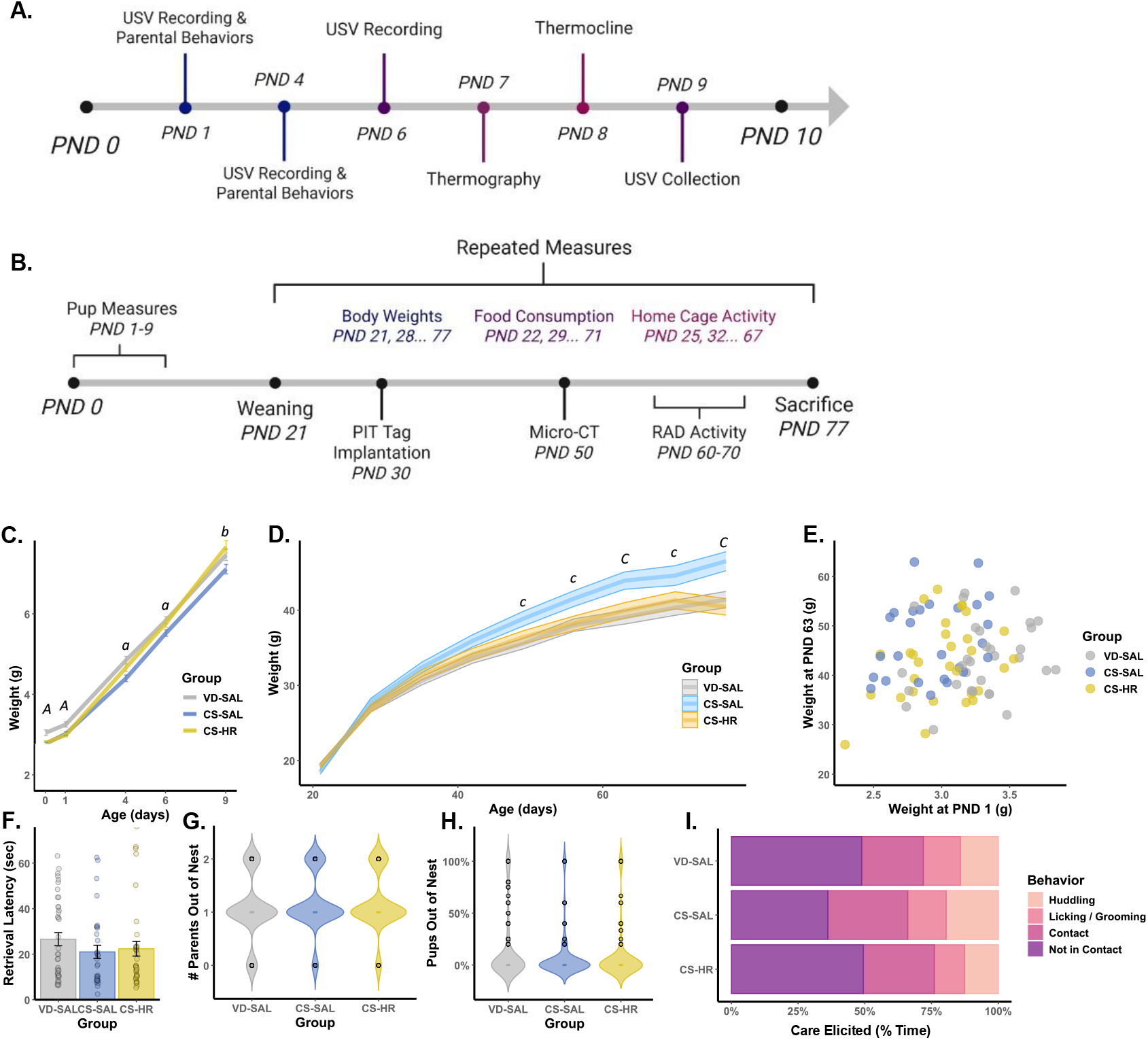
Delivery by CS leads to consistently heavier weights throughout development unless CS neonates are treated with hormone rescue (OXT and AVP, 1 mg/kg each, subcutaneous). Following birth, treatment, and cross-fostering, offspring were tested as pups (A) and throughout development (B) for behaviors relevant to thermoregulation, spontaneous locomotion, and food consumption to understand the source of metabolic dysfunction that led CS offspring to gain weight. C.) Pup body weights. A denotes VD-SAL pups weighed more than both CS-SAL and CS-HR; a denotes VD-SAL pups weighed more than CS-SAL; b denotes CS-HR pups weighed more than CS-SAL (n = 75-77, p < 0.05 for all denoted comparisons). D.) Offspring body weights post weaning. C denotes CS-SAL heavier than both VD-SAL and CS-HR; c denotes CS-SAL heavier than VD-SAL (n = 54-73, p < 0.05 for all denoted comparisons). E.) There was no relationship between birth weight and adult body weight. Pups from all groups experienced similar: retrieval latency (F), proportion of time parents spent in the nest (G), proportion of time spent outside the nest as pups (H), and elicited the same alloparental caregiving behaviors from an unrelated adult (I). Thus, CS-SAL subjects’ adult body weights appear unrelated to how they were cared for as pups.

### Parental Care, Pup Retrieval, and Alloparental Response

There were no group differences in retrieval latency or the proportion of pups successfully retrieved (Figure 1F). Likewise, there were no group differences in the amount of time either parents (Figure 1G) or pup (Figure 1H) spent outside the nest. When pups were given to adult males that had been previously vetted for alloparental responsiveness, these alloparents reacted similarly to pups from each group (Figure 1I). Taken together, these findings show that CS pups were equally effective as VD pups at eliciting care from adults.

### Ultrasonic Vocalizations

In our previous experiment [14], we observed conflicting effects of CS on the production of ultrasonic vocalizations, finding that CS can either increase or decrease ultrasonic vocalization production in voles (Figure 2).

**Figure 2.**
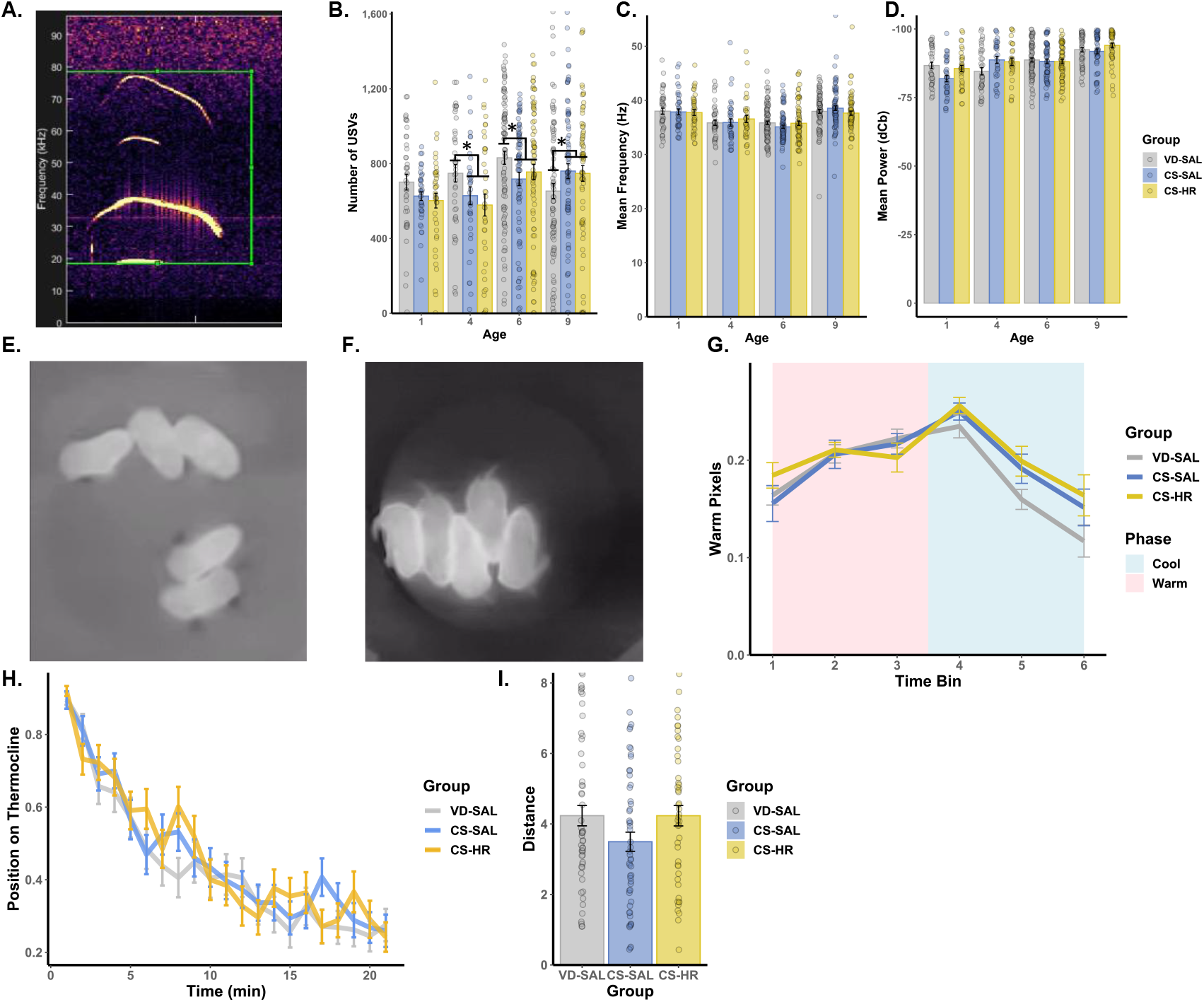
Pup behaviors. A.) Representative spectrogram of vole pup ultrasonic vocalization (USV). B.) Birth mode (rather than Group) impacted the number of USVs pups produced in an age-dependent manner, with VD pups initially vocalizing more on PNDs 4 and 6, then less on PND-9 (* denotes p < 0.05). There were no effects on mean frequency (C) or mean power (D). Representative thermographs of vole litters huddling under warm (30°, E) and cool (22°, F) conditions, for which there were no group differences in thermal emission (G). When pups were tested on a thermocline, there were likewise no group differences in distance traveled (H) or temperature preference (I).

### Thermoregulation as Pups

Having previously found that CS pups have lower surface temperatures in warm conditions and huddled less cohesively [14], we sought to replicate and extend these findings in 13 VD-SAL litters, 10 CS-SAL litters, and 8 CS-HR litters. However, there were no group effects on surface temperature of PND-7 pups (Figure 2G). On PND-8, pups were tested individually (n = 53 VD-SAL, 54 CS-SAL, and 46 CS-HR pups) on a thermocline where again there were no group effects on preferred temperature (Figure 2H) or movement (Figure 2I).

### Food consumption

There was no effect of group on food consumption (Figure 3A).

**Figure 3.**
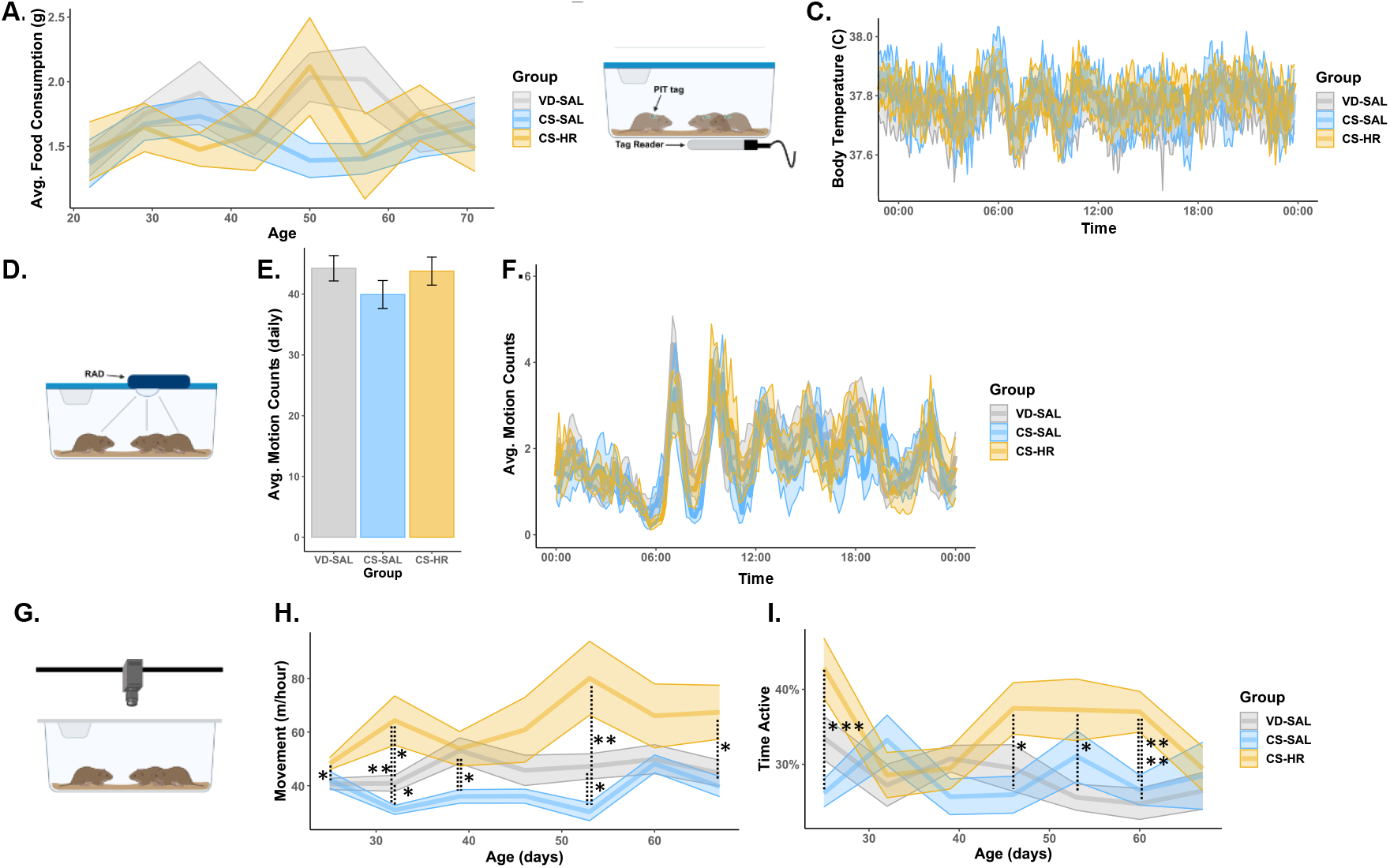
Energy balance. A.) There were no group differences in average food consumed. B.) Schematic of continuous body temperature recording. C.) There were no group differences in body temperature. D.) Schematic of home cage locomotor assessment by RAD. E.) There were no group differences in the proportion of time spent active across (E) or within (F) days. G.) Schematic of home cage locomotor assessment by video and idTracker. H.) CS-SAL subjects traveled less than VD-SAL and CS-HR throughout development, * p < 0.05, ** p < 0.01. I.) CS-HR subjects spent more time active than VD-SAL and CS-SAL throughout development, * p < 0.05, ** p < 0.01, *** p < 0.001.

### Body Temperature

Continuous measurement of body temperature via subcutaneously injected interscapular PIT tags (Figure 3B) revealed the expected ultradian rhythm typical of prairie voles, but did not detect any group differences in body temperature (Figure 3C)

### Home cage Locomotor Activity

Continuous detection of activity in the home cage using lid-mounted Rodent Activity Detectors (RADs, Figure 3D) also revealed the expected ultradian rhythm in motion, but did not detect any group differences in the proportion of time spent active (Figures 3E and F). Because the RAD only detects the presence or absence of motion, but not the degree of motion, we supplemented this measure with weekly 2-hour video sampling of behavior in the home cage (Figure 3G). Weekly video recording of the home cage revealed that CS-SAL generally moved less than either VD-SAL or CS-HR (Figure 3H). Thus, CS-SAL animals spent the same amount of time active but covered less distance during spontaneous activity in the home cage than VD-SAL counterparts, which could have contributed to lower overall energetic expenditure.

### Morphometry

At sacrifice, we collected measures of subjects’ overall weight, length, and the weights of several subcutaneous adipose deposits as well as that of the liver (Figure 4). There was an effect of Group on weight (F(2,69) = 10.91, p < 0.001), BMI (F(2,69) = 4.45, p < 0.015), and total adipose weight (BAT + WAT, F(2,72) = 3.937, p = 0.024) but not BAT or WAT individually, nor length. Post-hoc testing showed that CS-SAL had greater body weights at sacrifice than both VD-SAL and CS-HR (p < 0.01 for both comparisons) and greater BMI and total adipose than CS-HR (p < 0.03 for both comparisons).

**Figure 4.**
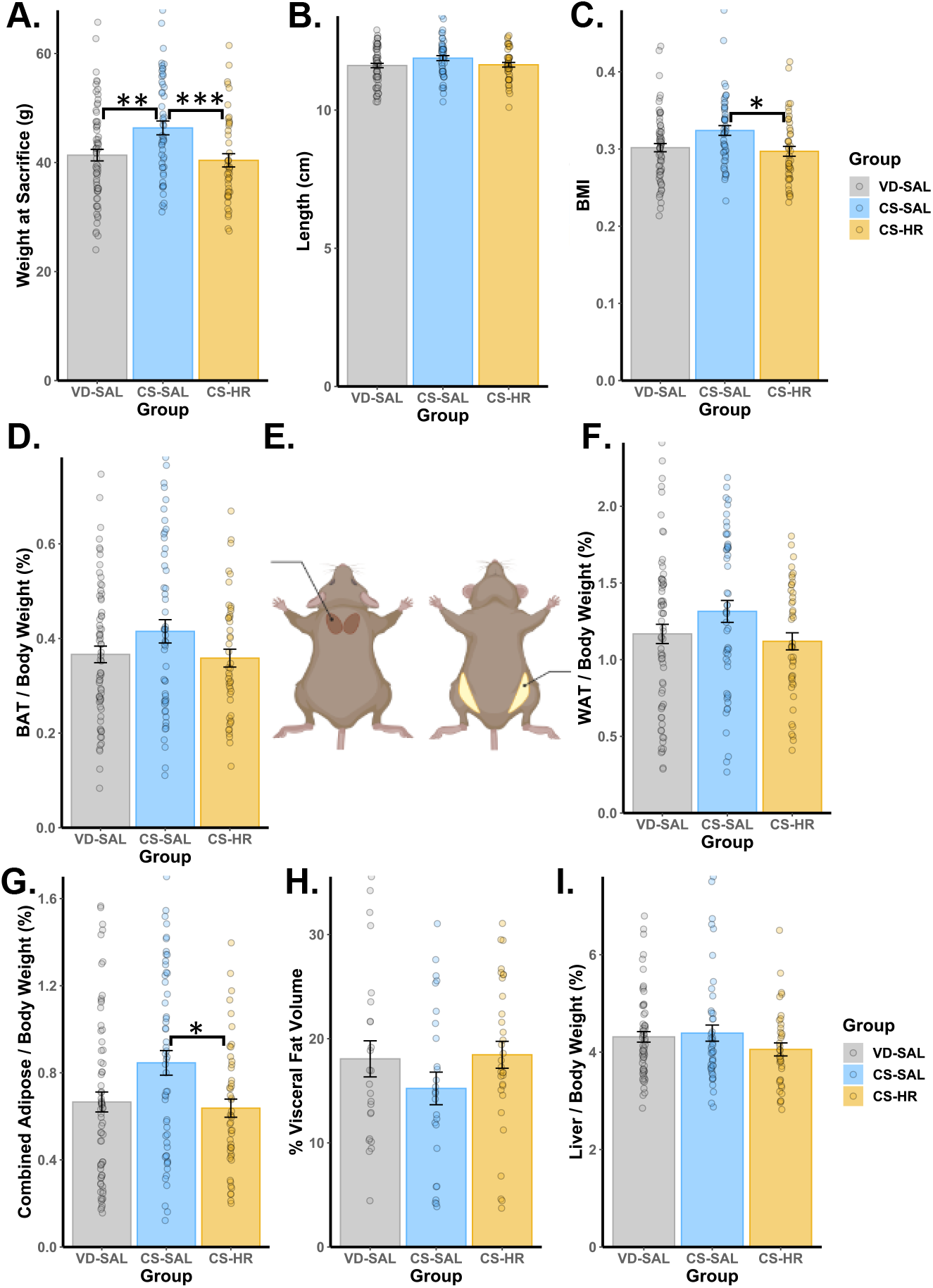
Morphometric results. A.) CS-SAL subjects exhibited greater body weights at sacrifice than either VD-SAL (** p < 0.01) or CS-HR (*** p < 0.001). B.) There were no group differences in body length. C.) CS-SAL subjects exhibited greater body mass index (BMI) than CS-HR (* p < 0.05). D.) There were no group differences in BAT weight. E.) Schematic of adipose depots. F.) There were no group differences in WAT weight. G.) CS-SAL subjects had greater total adipose depot weights than CS-HR (* p < 0.05). There were no group differences in either visceral adiposity (H) or liver weights (I).

### Brain Measures

At sacrifice, we collected brain tissue from subjects to be processed either for quantification of the OXT and AVP ligands (see: immunohistochemistry) or the OXT and AVP receptors (OTR and V1aR) and dopamine transporter (DAT; see autoradiography). OXT and AVP immunoreactivity was quantified in the paraventricular nucleus of the hypothalamus (PVN), one of the source nuclei for these neuropeptides (Figure 5). After observing the decreased spontaneous locomotor activity in the home cage, we also added brain tissue samples from a parallel cohort of voles, from which we immunohistochemically quantified tyrosine hydroxylase (TH), the rate-limiting enzyme in the synthesis of dopamine, across the anterior-posterior extent of the caudal caudate/putamen. Following that, DAT density was quantified across the anterior-posterior extent of the caudal caudate/putamen (Figure 6). OTR and V1aR density was quantified across an anterior-posterior extent of the forebrain ranging from the nucleus accumbens to the ventromedial hypothalamus (Supplemental Figures 1 and 2).

**Figure 5.**
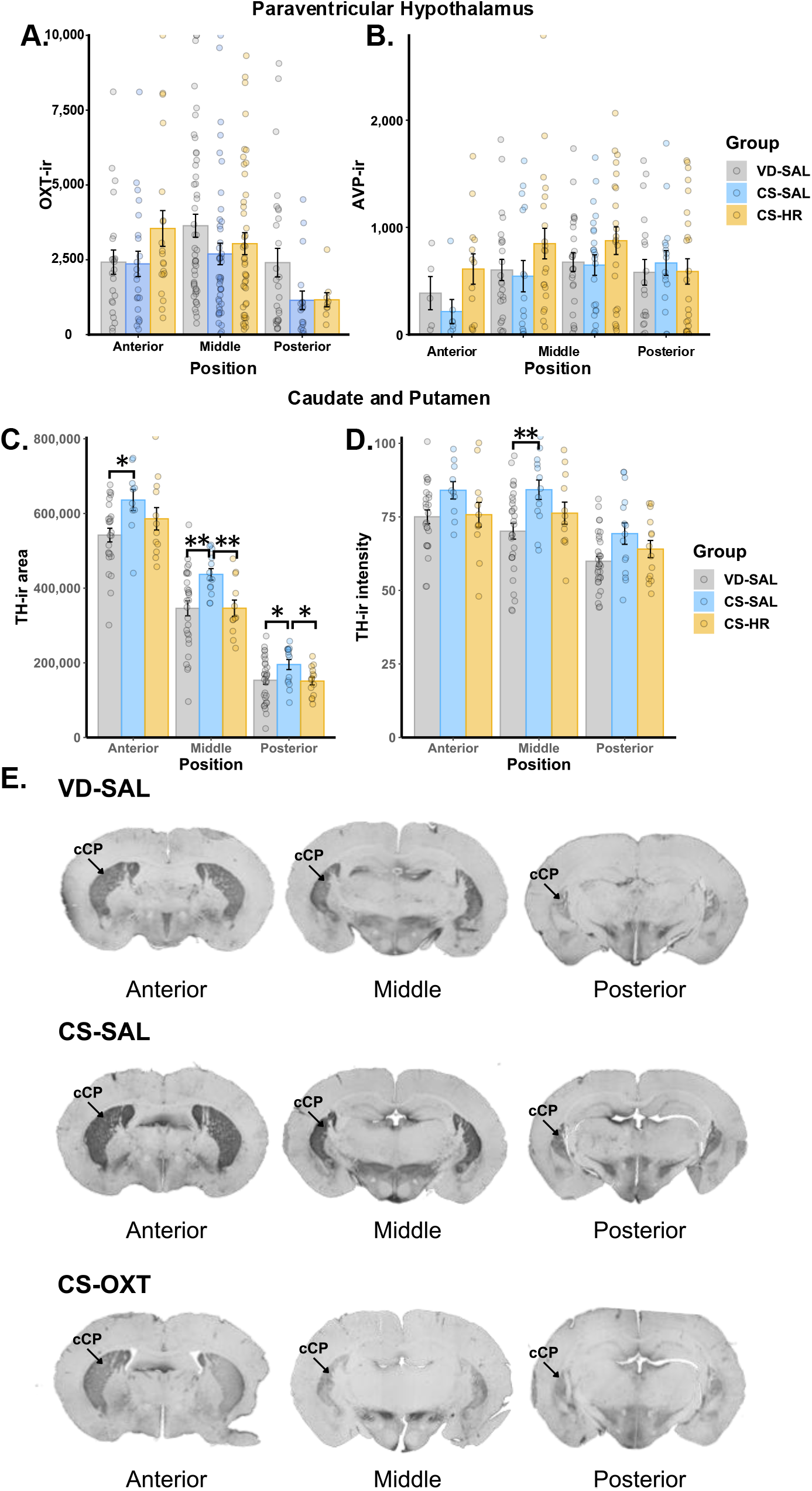
Immunohistochemistry throughout the brain. There were no group differences in either OXT-ir (A) or AVP-ir (B) within the PVN. C.) CS-SAL subjects had greater area of TH-ir across the cCP than VD-SAL or CS-HR (* p < 0.05, ** p < 0.01). D.) CS-SAL subjects had greater intensity of TH-ir than VD-SAL in the middle portion of the cCP (** p < 0.01). Representative photomicrographs of TH-ir in the cCP.

**Figure 6.**
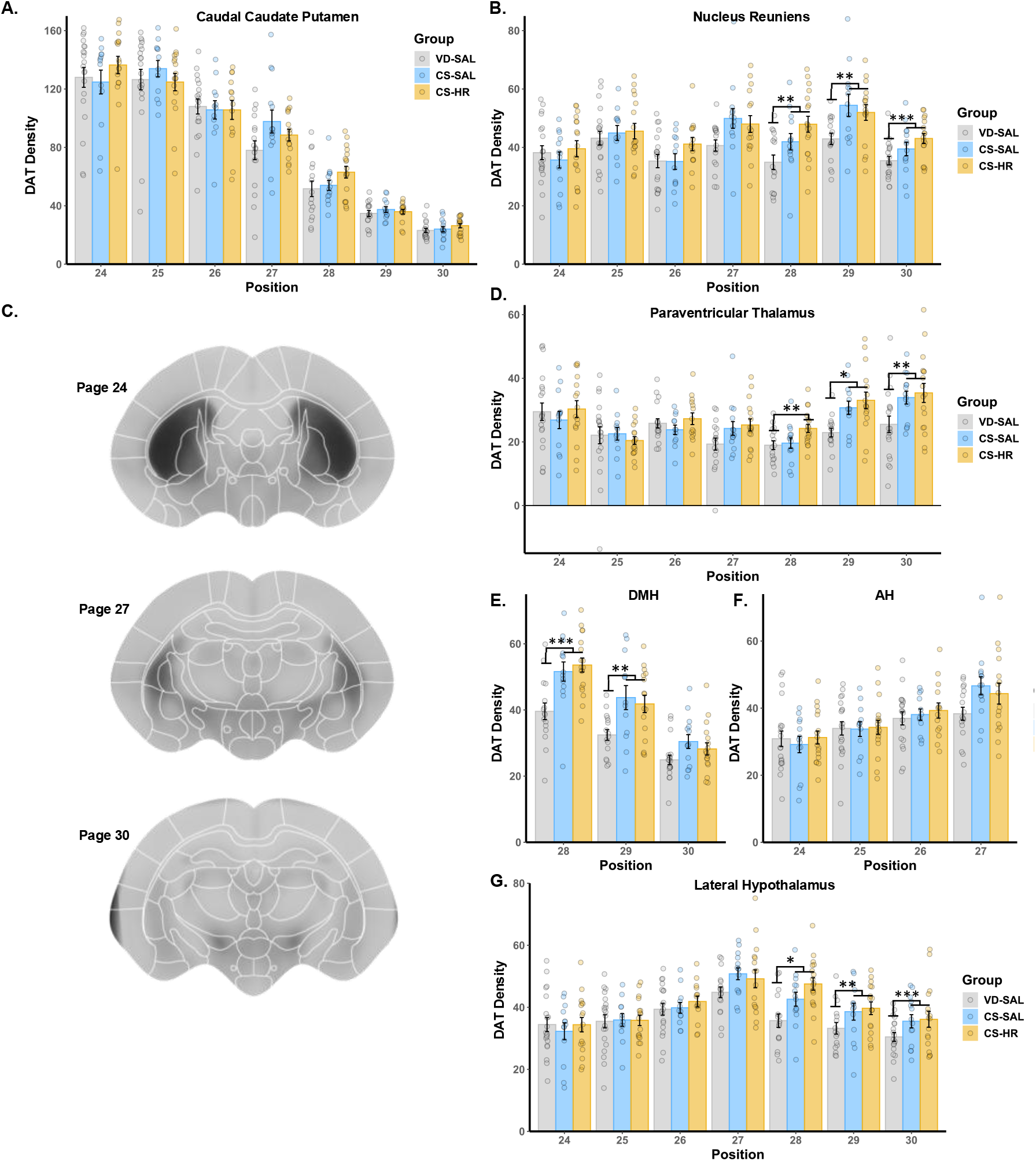
Dopamine transporter (DAT) autoradiography throughout the brain. A.) There were no group differences in DAT density throughout the CP. B). CS subjects had greater DAT than VD subjects in the posterior nucleus reuniens C.) Sample composite images of DAT autoradiography with atlas delineations overlaid. D.) CS subjects had greater DAT than VD subjects in the posterior paraventricular thalamus. E.) CS subjects had greater DAT than VD subjects in the dorsomedial hypothalamus (DMH). F.) There were no group differences in the anterior hypothalamus (AH). G.) CS subjects had greater DAT than VD subjects in the posterior lateral hypothalamus. Data are shown in 8-bit grayscale intensity relative to background. * p < 0.05, ** p < 0.01, *** p < 0.001.

### Immunohistochemistry

We observed no effects on either OXT or AVP immunoreactivity levels (area) across the entire anterior-posterior extent of the PVN (Figure 5A and 5B). In quantifying TH across the caudal caudate-putamen (cCP) in a separate cohort of animals, we measured both the area of expression and the intensity across the anterior-posterior extent of the caudal subdivision. We observed a consistent pattern of CS-SAL having greater area and signal intensity than both VD-SAL and CS-HR subjects (Figure 5C and 5D). Importantly, for this measure, CS-HR animals received only 0.1 mg/kg OXT, not AVP as in the other measures. We also retrospectively analyzed brains from a previous study [12] and found again that CS subjects had greater TH-ir intensity in the cCP (Supplemental figure 1).

### Autoradiography

We observed no effects of group on either OTR or V1aR density throughout any of the candidate regions (Anterior / Dorsomedial / Paraventricular / Ventromedial Hypothalamus, Caudate Putamen, or Globus Pallidus) or exploratory analyses (after correction for multiple comparisons) we examined (Supplemental Figures 2-3). Further results, including regional heatmaps and tables of exploratory results are presented in Supplemental Figures 2-3. We found no differences in DAT density in the cCP. Exploratory analyses of DAT density with corrections for multiple comparisons revealed a significant effect of group in the Reuniens Nucleus (RE), Paraventricular Thalamus (PVT), Dorsomedial Hypothalamus (DMH), Anterior Hypothalamus (AH), and Lateral Hypothalamus (LH), with CS-SAL and CS-HR subjects showing greater DAT density than VD-SAL subjects (Figure 6).

## Discussion

The primary finding of this study is that hormone rescue with an OXT and AVP cocktail can ameliorate CS-induced weight gain in voles that otherwise begins 4 weeks post-weaning. Our results eliminate several possible explanations for the weight gain seen in our CS delivered voles. There were no early life characteristics, other than birth mode and hormone exposure, that could explain adult body weight. Similarly, we were unable to explain adult body weights via differences in either food consumption or thermoregulation. Instead, locomotor differences were uncovered such that CS-SAL voles moved slower than VD-SAL, prior to the detection of body weight differences, which was reversed by hormone rescue with OXT and AVP. In a separate cohort, CS similarly led to greater TH-ir within the cCP which was also reversed by hormone rescue with OXT alone. These findings lead us to conclude that CS delivery leads to dysregulated striatal dopamine, which leads to diminished spontaneous locomotion in the home cage, which contributes to weight gain.

Alterations in dopaminergic systems have been a consistent finding in cesarean delivered offspring [19,24,25]. Such effects have been partially normalized by a subcutaneous injection of epinephrine in the neonate rat [26]. Social deficits have also been observed in CS delivered offspring and have been mitigated with subcutaneous injections of OXT either perinatally or during the first days of life [14,27]. Developmental programming via perinatal OXT and AVP has been demonstrated through natural, endogenous processes and through controlled, exogenous administration [11]. Vole offspring exposed to increased maternal OXT levels at birth show widespread changes in network-level organization, with robust effects on striatal functional connectivity and volume [28]. What’s more, adult rats given a single injection of OXT (5 mg/kg) on PND 1 have lower levels of DA metabolites in the striatum [29], further supporting that perinatal OXT manipulations can have lasting effects on DAergic neurotransmission and metabolism.

Thus, birth is a critical window for central nervous system development, during which perinatal hormone exposure significantly shapes DAergic organization.

The hypoactivity we observed in CS offspring mirrors those of prior open-field tests in mice, wherein CS offspring were observed to display less locomotion during 10-minute tests [27,30]. Gross motor deficits have also been noted in CS-delivered human children [31,32]. Later in development, CS-delivered children aged 7-10 are 44% less likely to have high levels of physical activity relative to VD counterparts [33]. This is the first study linking hypoactivity to weight gain in CS offspring.

In this study we observed that CS-SAL subjects had greater TH-ir area than VD-SAL and CS-HR subjects across the cCP, and higher TH-ir levels in the middle region of the cCP than VD-SAL subjects (Figure 5). These results are counterintuitive considering TH-ir area and levels were greater in the cCP and locomotor activity was decreased in CS-delivered subjects. This suggests the changes in TH seen here may be compensatory effects of some other dysfunction in striatal dopamine signaling.

The work featured several strengths of experimental design that bolster its conclusions.

Our cesarean section procedure does not utilize surgical anesthesia, so differences in the offspring cannot be attributed to anesthesia exposure. We also used universal cross-fostering, wherein CS and VD pups were both reared by non-biological parents, to exclude confounds of parental effects of CS. Whole litters were treated as subjects here, which excludes litter effects [34,35].The use of voles as subjects here also offers several advantages. First, voles tolerate room temperature housing well [36], without the metabolic [37,38], autonomic [12,39] and neuroendocrine [40] consequences faced by mice housed at room temperature. Voles are undomesticated, which means their OXT and AVP systems have not undergone artificial selection as in mice [41].

However, this study is also not without weakness. In this experiment, a mixed-chow was utilized that contained a combination of a vole diet and the relatively higher fat, standard mouse diet. Because these diets were presented in a heterogenous mixture, we have no way of knowing if one group preferentially consumed the calorically dense mouse chow. We did not find a difference in adiposity, despite CS animals weighing more, thus we cannot conclude the subjects became obese. While our home cage activity monitoring data suggest differences in spontaneous locomotor underlie the differences in body weight observed in our subjects, we cannot quantify the proportion of energy expenditure through movement with our current technology. While we speculate the brain differences observed in our subjects lead to greater weight gain, all brains were retrieved from PND-77 subjects, a timepoint well beyond when the weight differences began to emerge. Thus, we cannot determine whether brain differences drove weight increase or the reverse.

Given the increasing prevalence of CS deliveries globally, studies of potential adverse developmental consequences are essential. This need is especially apparent given the procedure’s association with obesity. Here, we showed that a simple, low risk injection of widely available hormones may be sufficient to prevent cesarean section induced weight gain.

## Supporting information

Supplemental Methods and Results

## Acknowledgments

The authors would like to relate their gratitude for the work of the Office of Laboratory Animal Medicine at the University of Delaware.

## Funding

This work was supported by:

National Institutes of Health grant P20GM103653 (WMK)

National Institutes of Health grant R01HD111737 (WMK, JEB)

This material was based upon work supported by the Office of Research and Development, Medical Research Service, Department of Veterans Affairs (VA). This work was also supported by the VA Merit Review Award 5 I01BX004102, from the United States (U.S.) Department of Veterans Affairs Biomedical Laboratory Research and Development Service grant to James Blevins.

The contents do not represent the views of the U.S. Department of Veterans Affairs or the United States Government.

