## Supplemental Methods and Results for "Oxytocin and Vasopressin at Birth Prevent Hypoactivity and Excess Weight Gain in Vole Offspring Delivered by Cesarean Section"

**This PDF file includes:**

Supporting text
Figures S1 to S3

**Supplemental Methods**

**Birth Conditions:** Doses were selected based on prior studies using higher concentrations (~2-10x greater (Bales and Perkeybile, 2012; Kenkel et al., 2023a, 2019)), with the goal of employing conservative, physiologically reasonable levels.

**Home Cage Parental Behavior:** Testing occurred following the removal of most of the nesting material (which was necessary to ensure visibility). A standardized area where the nest had been (10 x 10 cm) was designated as the nest area; if the parents relocated the majority of the litter to a new nest area, then a new 10 x 10 cm area at those coordinates became the nest area. Once each minute, the number of parents that had at least two paws within the boundaries of the nest area was recorded, as was the number of pups outside the nest area.

**Ultrasonic Vocalizations:** Ultrasonic recordings were captured using a Pettersson M500-384 microphone (Pettersson Elektronik AB Uppsala Sweden) and Audacity recording software (www.audacityteam.org). Recordings were conducted at 192000 Hz. Recordings were later quantified for the total number of vocalizations and total duration of vocalization using DeepSqueak (Coffey et al., 2019). The total number of vocalizations as well as the mean amplitude and frequency were calculated for each recording.

**Alloparental Response:** Approximately 30 minutes after the conclusion of pup retrieval testing on PNDs 1 and 4, two pups from each litter were randomly selected to be tested for their ability to elicit an alloparental response from an unrelated adult. Using a stock of six adult virgin males, pre-screened for alloparental tendencies, subject pups were removed from their home nest and placed into a clean, new cage along with one alloparental male for 20 minutes.

**Thermography:** Litters were placed on a thin foam pad within a 2L jacketed beaker measuring ~12.5 cm in diameter internally. The ambient temperature was initially set to 33° and maintained there for 30 minutes, at which time it was lowered to 22°. Once every minute, the litter was photographed simultaneously in the visual spectrum for behavior and in infrared to assess surface temperature (FLIR ONE Pro, Teledyne FLIR LLC) suspended 25 cm above the chamber floor. These minute-by-minute measures were averaged over 10-minute bins. The primary result consisted of the number of pixels at or above 34°. Alongside the infrared thermography, photographs were also captured once every minute to assess behavioral thermoregulation (i.e. huddling). Behavioral thermoregulation measures included the number of huddles and the total perimeter of all huddles, as we have reported previously (Kenkel et al., 2023a). Results were normalized to account for litter size, total litter body weight, and starting background temperature.

**Thermocline:** The thermocline consisted of a thin copper platform on which the temperature varied from ~20° to ~40°. The temperature range was confirmed prior to testing using infrared thermography (FLIR Inc.). This apparatus was constructed in-house and consisted on one end of an insulated cooler providing a heatsink of water into which the 20° end of the copper platform was placed, while on the other end, the copper platform was warmed to 40° by a commercial sous vide water heater (Anova Culinary Inc.) that continuously circulated warm water under the copper platform by means of a radiator from a 2001 Chevrolet Impala. Testing on the thermocline lasted 20 minutes and subjects' positions were scored once every 60 seconds.

**Body temperature:** On PND-30, a subset of subjects was lightly and briefly anesthetized with 1% isoflurane for 1-2 minutes while they were injected subcutaneously on the dorsum with a temperature sensor (BioTherm13 passive integrated transponder, BioMark Inc.). From PND-30 to PND-65 (mean = 44.3 days), these subjects were recorded for 1-7 days at a time. Recordings consisted of placing an antenna under the home cage and collecting body temperature measures once per minute when the subject was directly overhead the antenna (diameter ~18 cm). Thus, these recordings were experienced by the subjects as no different than standard housing. Measurement collection was automated using BioMark Device Manager software and then further processed in R. Having observed no effect of age on body temperature, we averaged temperature across the multiple days of the recording period to produce a within-subject average of the 24 hours of a typical day for each subject. These 24-hour consolidations were then used as the basis for statistical and graphical comparisons. A single subject (a VD-SAL male) was removed as an outlier for having a mean body temperature of 38.76° while the mean body temperature for all other subjects was 37.77° ± 0.27°.

**Home cage locomotor activity:** From PND-25 to PND-67, a subset of subjects were subjected to 2 hour recordings in the home cage every week. During these recordings, the cage top was replaced with a transparent plexiglass lid and the food and water removed. Behavior was recorded via overhead cameras and the total distance traveled was tracked via idTracker (Perez-Escudero et al., 2014). Sample sizes varied across weeks due to occasional technical issues, difficulties in subject identification, or logistical constraints that prevented video recording in some sessions.

From PND-60 through PND-77 (mean = 69.7 days), a subset of subjects' home cages was equipped with automated locomotor activity detectors. These devices, the Rodent Activity Detector (RAD) (Matikainen-Ankney et al., 2019), rest atop the cage lid and use infrared motion sensors to gauge activity below. When placed atop the cage lid, we ensured the sensor was on the opposite side of the cage from the nest so as to avoid observing motion within the nest. The most common issues we encountered were instances when subjects gaining access to and chewing the devices' wires, which stopped data collection, and instances when subjects knocked the devices over so that locomotor activity could not be detected. We therefore inspected each recording and automatically excised all instances of zero activity included in spans longer than 500 minutes (8.33 hours) of zero activity detected. The RAD continuously detects the presence / absence of movement in the home cage without disrupting subjects, whereas our home cage video recording provided more detailed quantification of distance traveled but only in 2-hour observations that entailed relocating the home cage and removing its lid.

Locomotor activity was normalized to the number of voles in the cage. Results were analyzed both in terms of the proportion of time any activity was detected as well as the average motion events observed during a given minute. One cage of CS-HR females' apparent high rate of activity marked it as an outlier ($p < 0.001$ in Grubbs' test) and so it was removed from the analysis. Because of the small sample sizes when considering sex and treatment group concurrently, we decided to collapse across sex. Having observed no effect of age on locomotor activity, we averaged locomotion across the multiple days of the recording period to produce a within-cage average of the 24 hours of a typical day for each cage. These 24-hour consolidations were then used as the basis for statistical and graphical comparisons.

**Micro-CT:** On PND-50, a subset of subjects were lightly anesthetized with 2% isoflurane and 3D-imaged with a Brüker SkyScan 1276 (Kontich, Belgium) at a nominal resolution of 21.2 microns using a 1mm thick aluminum filter and an applied x-ray tube voltage of 60 kV. Camera pixel binning of 2x2 was applied. The scan orbit was 180 degrees with a rotation step of 0.9 degrees. Imaging was restricted to the abdomen defined as from the lowest rib to the top of the pelvis. Reconstruction was carried out with a modified Feldkamp algorithm using the Brüker SkyScanTM NRecon software and NReconServer (version 1.7.4.2). Beam hardening correction was applied.

**Tissue Collection:** Vole length and final body weight were recorded. Length of vole was measured from nose to base of tail. After cervical dislocation, necropsy was conducted to collect various tissues for analysis and preservation. Briefly, brains were either flash frozen on dry ice for autoradiography or fixed for immunohistochemistry. All tissue samples were weighed immediately following excision and weights were summed bilaterally.

**Immunohistochemistry:** Brains were immersion-fixed in 4% paraformaldehyde for 24 hours followed by at least 24 hours in 20% sucrose before being sectioned at 30μm. Immunohistochemical staining was carried out in free floating tissue according to the following procedure. First, brains were rinsed three times for 5 minutes each in KPBS, followed by incubation in 1% $H_2O_2$ in KPBS for 30 minutes and another three 5-minute washes in KPBS. Slices were then incubated in primary antibodies (either AVP at 1:16,000 dilution, OXT at 1:16,000, or tyrosine hydroxylase (TH) at 1:3500 MilliporeSigma antibodies # AB1565, MAB5296, and IHCR10056, respectively) in KPBS plus 0.4% Triton-X for one hour at room temperature and another 60 hours at 4°. This was followed by three more 5-minute KPBS washes and 30 minutes in KPBS + triton-X and 1% normal horse serum (OXT and AVP) or goat serum (TH). Tissue was then incubated in KPBS + triton-X plus biotinylated horse anti-mouse/rabbit secondary antibody (OXT and AVP) or KPBS + triton-X plus biotinylated goat anti-mouse/rabbit secondary antibody (TH) at 1:600 (Vector Laboratories, antibody # BA92001.5). Tissue was then washed five times and incubated in A/B solution followed by three more washes in KPBS and three more in Na-acetate before chromogen precipitation brought about by Ni-DAB. After further washing, tissue was mounted and cover-slipped.

**Histological Quantification:** Images from brain tissue were collected using a EVOS M7000 Microscope Imaging System at 4x magnification, with entire slides digitally stitched together to form single images. Images were then quantified using ImageJ (Schneider et al., 2012) to assess the immunoreactivity (ir). To account for non-specific binding and differences in image brightness, background values were taken from signal-negative areas, and then the mean ROI pixel intensity was subtracted from the mean background pixel intensity. OXT-ir and AVP-ir were assessed using a standardized region of interest (ROI) along the anterior-posterior extent of the PVN (pages 24-27 in our vole brain atlas), and TH-ir was assessed using manually set ROIs over the left and right cCP (caudal caudate-putamen (cCP, pages 24-30) and SN (pages 32-34). TH-ir values were averaged across the left and right ROIscCP for analysis. Alternate slices from each individual were used for OXT and AVP. For OXT-ir and AVP-ir, signal strength was maximal so as to capture cell fibers, so we analyzed the area of staining. For TH-ir, signal strength varied, so we analyzed both the area and intensity of staining. For anatomical reference, we used the Prairie Vole Brain Atlas and the Allen Brain Atlas (Allen Institute for Brain Science, 2011) and included anterior-posterior position as a covariate in the analyses.

**Autoradiography:** Flash frozen brain tissue was sectioned at 20μm at -15-20°C on a cryostat into six adjacent series and were thaw-mounted to SuperFrost Plus slides. Slides were stored at -80°C until use in autoradiography assays. Receptor autoradiography for oxytocin receptor (OTR) and arginine vasopressin receptor type 1a (V1aR) was performed with methods validated in prairie voles (Rogers et al., 2021) and outlined briefly here. A full anterior to posterior series from every subject was removed from −80°C storage and brought up to room temperature. The mounted brain tissue was then lightly fixed with 0.1% buffered paraformaldehyde (pH 7.4) for 2 minutes, washed twice for 10 minutes in 50 mM Tris buffer (room temperature, pH 7.4). Slides designated for analysis of OTR were then incubated for 1-hour in tracer buffer (50 mM Tris buffer + 10 mM MgCl, 0.1% BSA, pH 7.4) containing 50 pM of $^{125}$I-Ornithine Vasotocin Analog ($^{125}$I-OVTA; Vasotocin, d(CH2)5[Tyr(Me)2, Thr4, Orn8, [$^{125}$I]Tyr9-NH2]; Revvity). Slides designated for analysis of V1aR were incubated for 1-hour in tracer buffer containing 50 pM of $^{125}$I-Linear Vasopressin 1a Receptor Antagonist ($^{125}$I-LVA; [$^{125}$I]-Phenylacetyl-D-Tyr(Me)-Phe-Gln-Asn-Arg-Pro-ArgTyr-NH2; Revvity). Following incubation, slides were washed twice in 4°C 50 mM Tris base with 10 mM MgCl (pH 7.4) for 20 minutes total, washed again in room temperature 50 mM Tris buffer with MgCl for 30 minutes with agitation, briefly dipped in deionized water, and then left to air dry. The slides were then exposed to UltraCruz Blue autoradiography film (Santa Cruz Biotechnology) for 7 days prior to development and analysis. A set of $^{125}$I standard slides (American Radiolabeled Chemicals, Inc) was also incubated on the film for creation of a standard curve in the analysis.

Dopamine transporter (DAT) autoradiography was conducted based on previously published studies in rodents (Fricks-Gleason et al., 2016; Murthy et al., 2008). To our knowledge, this is the first assessment of DAT binding in the prairie vole. One complete series of sections adjacent to those used in OTR and V1aR autoradiography was removed from −80°C storage and brought up to room temperature. Slides were lightly fixed in 0.1% buffered paraformaldehyde (pH

7.4) for 2 minutes and washed twice for 10 min each in 50 mM Tris buffer + 120 mM NaCl (room temperature, pH 7.4). Slides were then incubated for 2 hours in this same buffer (50 mM Tris buffer + 120 mM NaCl) containing 40pm $^{125}$I-RTI (Revvity) + 100 nM fluoxetine to block the known affinity of RTI for the serotonin transporter. This approach reliably reveals selective DAT binding. After incubation, slides were washed twice for 10 min each in cold 50 mM Tris buffer + 120 mM NaCl (4°C), dipped in deionized water, and air dried. The slides were then exposed to UltraCruz Blue autoradiography film (Santa Cruz Biotechnology) for 7 days and developed. Because the signal on the film was saturated, we waited one month for the signal to decay slightly before re-exposing the slides to film for 7 days to generate usable autoradiograms for quantitative analysis. A set of $^{125}$I standard slides was also incubated on the film for creation of a standard curve in the analysis.

**Autoradiographic Quantification:** Images of slides were digitally scanned at 1200dpi. Three images of each subject, matched for anterior-posterior position, were then registered to one another using the BigWarp plugin for ImageJ (Schneider et al., 2012). A measure of non-specific binding (NSB) was also taken for each section from a region where minimal binding is detected. The NSB value was subtracted from the binding value for each section and a mean was then calculated for each section. Optical density was converted into decays per minute (DPM) using the $^{125}$I standards. DPM were then measured within each atlas-defined ROI using custom designed Matlab code and the prairie vole brain atlas we originally developed for magnetic resonance imaging (Yee et al., 2016). Results were calculated as regional averages of DPM (which corresponds to receptor density). Images were then consolidated into group composites for the purpose of visualizing group differences. Heat maps were generated for each group-by-group comparison by subtracting one group composite from another (Supplemental Figures 2 and 3). We selected six regions as target ROIs (Anterior / Dorsomedial / Paraventricular / Ventromedial Hypothalamus, Caudate Putamen, or Globus Pallidus), which were analyzed individually. All remaining brain regions with relevant levels of receptor density were analyzed in a separate exploratory analysis. Analyses of autoradiography yielded 20 VD-SAL, 14 CS-SAL, and 18 CS-HR subjects.

**Supplemental Results**

**Weights Across Development:** In total, 17 VD-SAL litters, 15 CS-SAL litters, and 18 CS-HR litters were generated, yielding 75, 67, and 77 pups respectively. On PND-0, VD-SAL litters were heavier, weighing 3.05 ± 0.07g at discovery, while CS-SAL litters weighed 2.78 ± 0.03g, and CS-HR litters weighed 2.79 ± 0.05g at delivery, which resulted in a significant effect of group ($F(2,47) = 7.4$, $p < 0.002$). Preweaning weights showed an interaction between age and group ($F(2,581.45) = 6.87$, $p < 0.002$, Figure 1C), such that at VD-SAL pups weighed more than either CS-SAL or CS-HR on PND-1 ($p = 0.007$ and $p = 0.02$, respectively). Body weights were recorded from PND-21 to PND-91 from 15 VD-SAL litters, 11 CS-SAL litters, and 13 CS-HR litters, yielding 73, 56, and 54 individuals respectively. There were significant interactions between Sex and Age (as expected) and Group and Age ($F(2,534,45) = 21.56$, $p < 0.001$, Figure 1D), such that CS-SAL animals were consistently heavier than VD-SAL animals starting on PND-49 and heavier also than CS-HR animals on PND-63 and 77 ($p < 0.05$ for all comparisons).

**Ultrasonic Vocalizations:** In our two previous experiments, we observed conflicting effects of CS on the production of ultrasonic vocalizations, finding that CS can either increase or decrease ultrasonic vocalization production in voles (Kenkel et al., 2023a).

For ultrasonic vocalizations in particular, we observed an effect of birth mode (CS vs VD) rather than group, thus we collapsed between CS-SAL and CS-HR. Specifically, we observed an interaction between birth mode and age on the number of USVs produced ($F(1,658) = 9.67$, $p < 0.002$, Figure 2), such that on PNDs 4 and 6, CS pups vocalized fewer times than VD counterparts ($p = 0.018$ and 0.033, respectively), while on PND 9, CS pups vocalized more than VD counterparts ($p = 0.04$). There were no differences between CS-SAL and CS-HR pups and there were no group effects on mean amplitude or mean frequency of USVs.

**Thermoregulation as Pups:** Thermoregulation is a key adaptation to extrauterine life that is regulated by OXT, which we have previously observed to be acutely impaired in CS neonates (Kenkel et al., 2023a). Pups from all groups increased surface temperature as time continued in the 33° condition and shed warmth as once the ambient temperature dropped to 22°. There were also no group effects on behavioral thermoregulation, with litters from all conditions consolidating their huddling when temperature dropped to 22°. On PND-8, pups were tested individually (n = 53 VD-SAL, 54 CS-SAL, and 46 CS-HR pups) on a thermocline 90 cm long, where temperature ranged from 23° to 39°. Pups from all groups moved toward the warmer end of the thermocline at the same rate and while CS-SAL pups appeared to move less overall, this did not reach statistical significance (Figure 2I).

**Home cage Locomotor Activity:** Weekly video recording of the home cage revealed an age by group interaction (F(2,946.2) = 4.07, p = 0.017, Figure 3H), with CS-SAL generally moving less than either VD-SAL or CS-HR, such that on PND-25, VD-SAL moved less than CS-HR (p = 0.039); on PND-32, CS-SAL moved less than VD-SAL (p = 0.013) and CS-HR (p = 0.002), while CS-HR moved more than VD-SAL (p = 0.048); on PND-39, CS-SAL moved less than either VD-SAL (p = 0.006) or CS-HR (p = 0.033); on PND-53, CS-SAL again moved less than either VD-SAL (p = 0.011) or CS-HR (p = 0.003); and on PND-67, CS-SAL moved less than CS-HR (p = 0.035). To corroborate the findings from the RADs, we also calculated the time spent active during the weekly video recordings, and found that CS-HR subjects were more active than CS-SAL animals on PNDs 25, 46, and 60 and CS-HR were more active than VD-SAL on PNDs 53 and 60 (p < 0.05 for all comparisons). Thus, CS-SAL animals spent the same amount of time active but covered less distance during spontaneous activity in the home cage than VD-SAL counterparts, which could have contributed to lower overall energetic expenditure.

**Morphometry:** On PND-50, subjects underwent micro-CT scans of the abdomen (from the posterior rib to the pelvis). Given that terminal outcomes were measured at a single time point and exhibited substantial litter-level variance, we analyzed terminal measures (body length and adipose depots) using litter–sex averages. Body length was included as a covariate in the analyses of adipose depots and weight at sacrifice.

**Immunohistochemistry:** For TH-ir area, we observed a significant main effect of group (F(2,56.47) = 5.38, p < 0.01) and region (F(2,347.64) = 692.43, p < 0.001), and an interaction between region and sex (F(2,347.64) = 3.96, p = 0.02). Post-hoc analyses showed that CS-SAL subjects had a greater TH-ir area than VD-SAL subjects in the anterior (*d* = -1.028, p =0.033), middle (*d* = -0.95, p = 0.003), and posterior (*d* = -0.73, p = 0.05) cCP, and a greater area than CS-HR subjects in the middle (*d* = 1.38, p = 0.008) and posterior (*d* = 0.96, p =0.038) cCP (Figure 5C). For mean intensity, there was a significant main effect of group (F(2,56.47) = 4.33, p = 0.018) and region (F(2,345.92) = 52.41, p < 0.001). Post-hoc analyses revealed that CS-SAL subjects had higher TH-ir intensity in the middle cCP than VD-SAL subjects (Figure 5D; *d* = -1.02, p =0.007). We also retrospectively analyzed brains from a previous study (Kenkel et al., 2024) and found again that CS subjects had greater TH-ir intensity in the cCP (Supplemental figure 1). There was a significant main effect of group ((F(1, 21.074) = 5.041, p = 0.036) and region (F(2, 317.37) = 11.26, p < 0.001), and a significant interaction between group and region (F(2, 317.24) = 3.27, p = 0.039). Post-hoc analysis revealed that CS subjects had a greater signal intensity in the anterior (d = -0.61, p = 0.003) and posterior (d = -0.45, p < 0.0001) cCP than VD subjects. For area, there was no significant effect of group. One outlying data point from the VD-SAL males was removed from the anterior substantia nigra (SN). There was a significant main effect of page ($\chi^2(2)$ = 223.63, p < 0.001), and a significant interaction between group and page ($\chi^2(4)$ = 10.039, p = 0.04) on TH-ir in the SN. Post-hoc analyses revealed that CS-HR subjects have greater TH pixel intensity compared to CS-SAL subjects in the anterior SN (*d* = 1.55, p = 0.022; data not shown).

**Autoradiography:** Exploratory analyses of DAT density with corrections for multiple comparisons revealed a significant effect of group in the Reuniens Nucleus (RE; F(2, 754.37) = 4.94, p = 0.047), Paraventricular Thalamus (PVT; F(2,753.27) = 13.44, p < 0.001), Dorsomedial

Hypothalamus (DMH; $F(2,300.029) = 6.59$, $p = 0.023$), Anterior Hypothalamus (AH; $F(2,392.8) = 5.12$ , $p = 0.046$), and Lateral Hypothalamus (LH; $F(2, 752.52) = 4.71$, $p = 0.048$), with CS-SAL and CS-HR subjects showing greater DAT density than VD-SAL subjects. We observed a significant interaction between group and position in the RE ($F(2, 733.57) = 6.67$, $p = 0.023$), PVT ($F(2, 718.79) = 16.047$ , $p < 0.001$), DMH ($F(2, 297.98) = 5.82$, $p = 0.032$), AH ($F(2, 386.76) = 5.51$, $p = 0.036$), and LH ($F(2,733.33) = 5.82$, $p = 0.032$). Post-hoc analyses revealed that CS-SAL subjects had greater DAT density than VD-SAL subjects in the RE on positions 28 ($d = 0.14$, $p = 0.0086$), 29 ($d = 0.15$, $p = 0.002$), 30 ($d = 0.16$, $p < 0.001$), PVT on positions 29 ($d = 0.32$, $p = 0.026$) and 30 ($d = 0.35$, $p = 0.005$), DMH on positions 28 ($d = -0.04$, $p < 0.001$) and 29 ($d = -0.085$, $p = 0.0021$), and LH on positions 28 ($d = 0.086$, $p = 0.048$), 29 ($d = 0.104$, $p = 0.015$), and 30 ($d = 0.12$, $p = 0.0068$). CS-HR subjects had a greater DAT density than VD-SAL subjects in the RE on positions 27 ($d = -0.26$, $p =0.0056$), 28 ($d = -0.39$, $p < 0,001$), 29 ($d = -0.52$, $p < 0.001$), and 30 ($d = -0.65$, $p < 0.001$), the PVT on positions 28 ($d = -0.15$, $p = 0.0058$), 29 ($d = -0.32$, $p < 0.001$), and 30 ($d = -0.47$, $p < 0.001$), and the DMH on positions 28 ($d = -1.48$, $p < 0.001$) and 29 ($d = -1.085$, $p = 0.0021$), and LH on positions 28 ($d = -0.3$, $p = 0.011$), 29 ($d = -0.4$, $p = 0.002$), and 30 ($d = -0.5$, $p < 0.001$).

### Figures

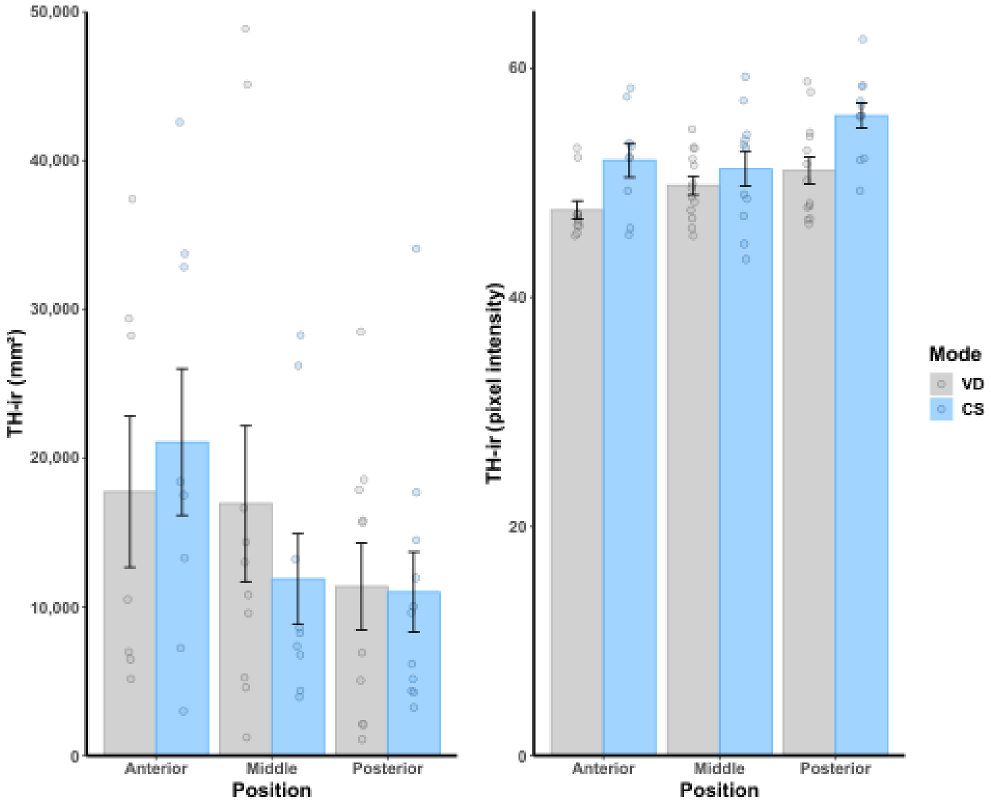


**Fig. S1.** TH-ir in the cCP. Results from a separate cohort, the subjects from [32]. (A) There was no effect on the total area of TH-ir in the cCP. (B) There were main effects of group and region, and a significant interaction between group and region, such that CS subjects had a greater signal intensity in the anterior (d = -0.61, ** p = 0.003) and posterior (d = -0.45, *** p < 0.0001) cCP than VD subjects.

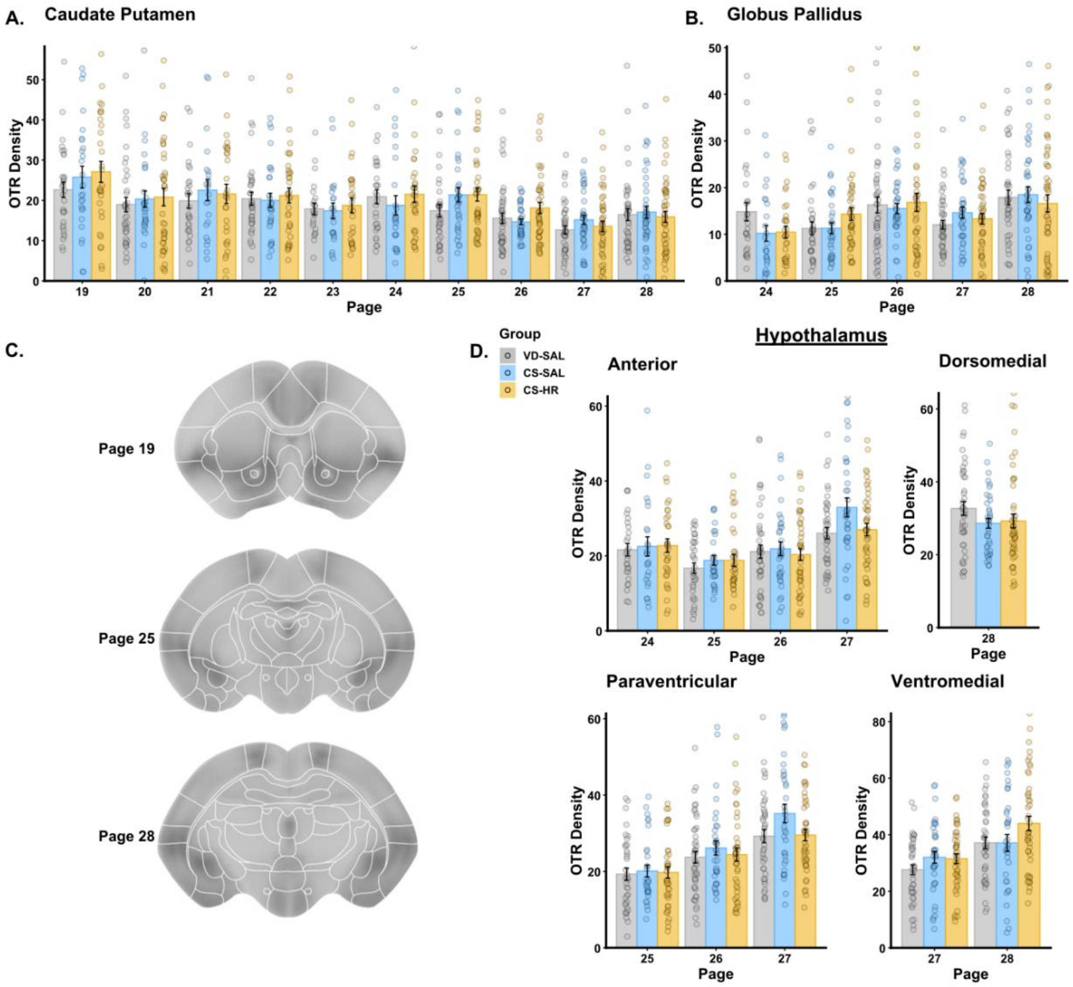


**Fig. S2.** OTR autoradiography throughout the brain. A.) There were no group differences in OTR density throughout the CP or in the globus pallidus (B). C.) Sample composite images of OTR autoradiography with atlas delineations overlaid. D.) There were no group differences in OTR density throughout the hypothalamus. Data are shown in 8-bit grayscale intensity relative to background.

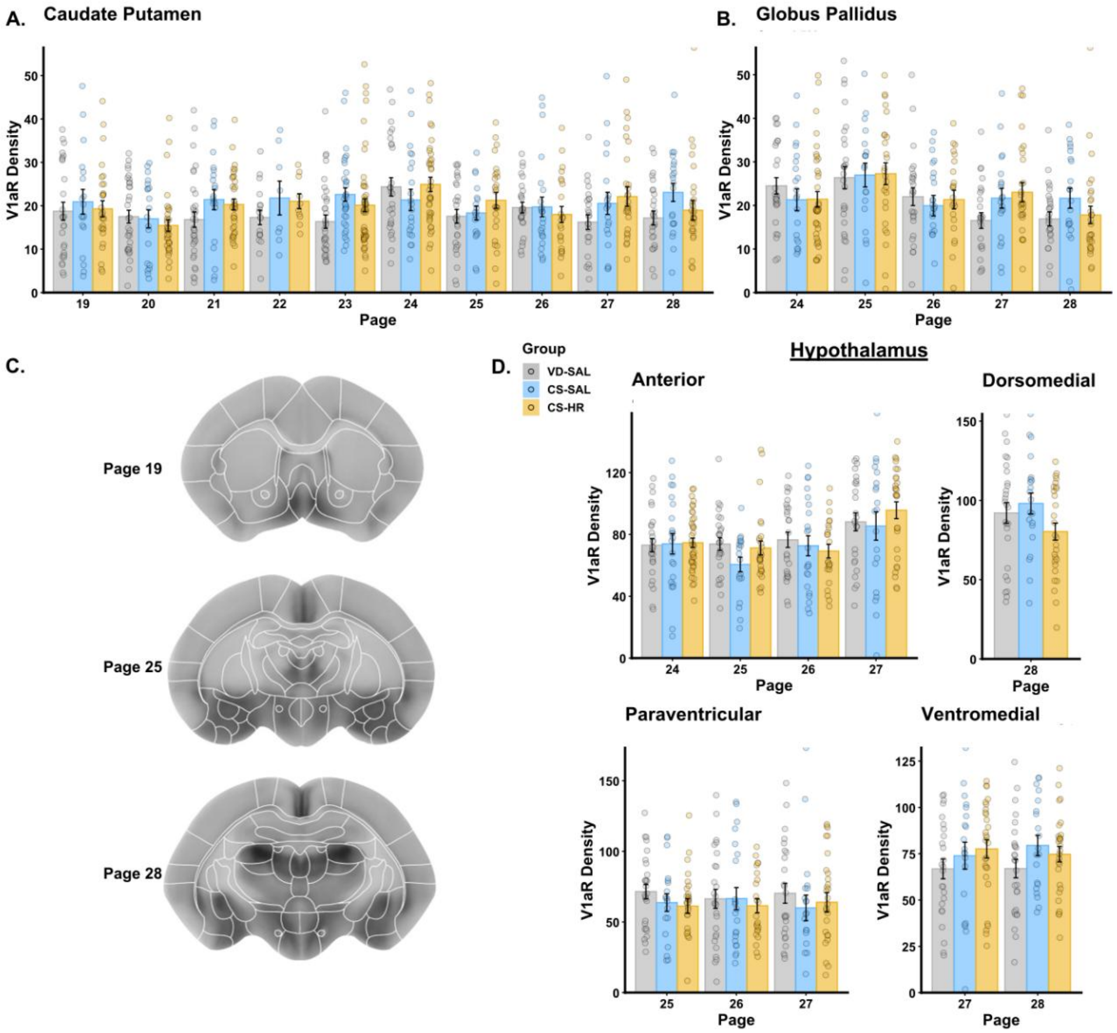


**Fig. S3.** V1aR autoradiography throughout the brain. A.) There were no group differences in V1aR density throughout the CP or in the globus pallidus (B). C.) Sample composite images of V1aR autoradiography with atlas delineations overlaid. D.) There were no group differences in V1aR density throughout the hypothalamus. Data are shown in 8-bit grayscale intensity relative to background.
